# An Open-Source Rodent Chronic EEG Array System with High Density MXene-Based Skull Surface Electrodes

**DOI:** 10.1101/2022.12.28.522126

**Authors:** Li Ding, Aashvi Patel, Sneha Shankar, Nicolette Driscoll, Chengwen Zhou, Tonia S Rex, Flavia Vitale, Martin J. Gallagher

**Affiliations:** Department of Neurology, Vanderbilt University School of Medicine, Nashville, TN 37232, USA; Department of Bioengineering, Center for Neuroengineering & Therapeutics, Department of Neurology, University of Pennsylvania, Philadelphia, PA, 19104 USA; Department of Ophthalmology & Visual Sciences, Vanderbilt University School of Medicine, Nashville, TN 37232, USA; Center for Neurotrauma, Neurodegeneration, and Restoration, Corporal Michael J. Crescenz Veterans Affairs Medical Center, Philadelphia, PA, 19104 USA; Tennessee Valley Health System, Department of Veteran’s Affairs, Nashville, TN 37212, USA

## Abstract

Electroencephalography (EEG) is an indispensable tool in epilepsy, sleep, and behavioral research. In rodents, EEG recordings are typically performed with metal electrodes that traverse the skull into the epidural space. In addition to requiring a major surgery, this intracranial EEG technique is difficult to perform for more than a few electrodes, is time-intensive, and confounds experiments studying traumatic brain injury. Here, we describe an open-source cost-effective refinement of this technique for chronic mouse EEG recording. Our alternative two channel (EEG2) and sixteen channel high-density EEG (HdEEG) arrays use electrodes made of the novel, flexible 2D nanomaterial titanium carbide (Ti_3_C_2_T_*x*_) MXene. The MXene electrodes are placed on the surface of the intact skull and establish electrical connection without conductive gel or paste. Fabrication and implantation times of MXene EEG electrodes are significantly shorter than the standard approach and recorded resting baseline and epileptiform EEG waveforms are similar to those obtained with traditional epidural electrodes. Applying HdEEG to a mild traumatic brain injury (mTBI) model in mice of both sexes revealed that mTBI significantly altered awake resting state spectral density with a spatiospectral region of interest in the β spectral band (12-30 Hz) in the central and posterior regions. These findings indicate that fabrication of MXene electrode arrays is a cost effective, efficient technology for multichannel EEG recording in rodents that obviates the need for skull-penetrating surgery. Moreover, increased β spectral power may contribute to the development of early post-mTBI seizures.

**Significance Statement:** Electroencephalography (EEG) is a critical technique used to study neurological activity in rodents. Commonly used EEG procedures require time-consuming skull-penetrating surgeries that may confound the experiments. Here we provide a cost-effective solution for obtaining two channel (EEG2) and high-density EEG (HdEEG) recordings on the skull surface thus avoiding major surgery. We compared this HdEEG system to traditional EEG recordings and then used it to determine the effects of mild traumatic brain injury on awake resting state spectral density. This novel open-source EEG system will contribute the electrophysiological characterization of rodent behaviors and seizure activity.

## 1. Introduction

Electroencephalography (EEG) is an indispensable tool in epilepsy, sleep, and behavioral research. The most common role of EEG in rodent experiments is to quantify seizures and stage sleep (del Campo et al., 2009) and thus only needs to record signals from a limited number of brain regions (often two channels, EEG2). However, experiments using high density EEG (HdEEG, 16-32 channels) are necessary for experiments designed to localize neurophysiological activity within the rodent brain with high resolution and accuracy.

Human EEG electrodes are metal cups filled with an electrolyte gel that establishes electrical contact with the scalp and must be repeatedly replaced during extended recordings due to gel drying (Searle and Kirkup, 2000). In addition to electrolyte drying, gel-filled cup electrodes cannot be used for chronic rodent EEG recordings due to the difficulty in attaching them to the scalp/skull and the propensity of the electrolyte to bridge adjacent electrodes on their small skulls. Therefore, current rodent EEG studies typically use 0.1-1.0 mm diameter metal electrodes inserted via cranial burr holes into the epidural space to establish electrode/electrolyte contact and to secure them for chronic recording. Drilling cranial burr holes is a major surgery that could potentially alter physiology. Skull penetrating surgeries are difficult in young rodents with thin skulls and they confound experiments studying mild traumatic brain injuries (mTBI) which are not associated with skull fracture. Epidural HdEEG experiments done with homemade arrays involve laborious fabrication and implantation procedures (Wasilczuk et al., 2016; Ding et al., 2019). While HdEEG experiments can also be performed using commercially available ultrathin 16-32 channel electrode arrays that lie on the skull surface (Choi et al., 2010; Jonak et al., 2018, 2020) they still require the placement of three epidural anchoring screws and their cost is prohibitive. There is a need for inexpensive, easily fabricated EEG2 and HdEEG arrays that can be implanted without skull injury.

In this open-source EEG2/HdEEG method, investigators design thin flexible printed circuit boards (PCBs) with electrode pads overlying cortical regions of interest. The electrode pads are then functionalized with soft contacts made with titanium carbide (Ti_3_C_2_T_*x*_) MXene, a biocompatible (Dai et al., 2017), flexible 2D nanomaterial that provides electrical contact between the electrode pad and the skull without the need of electrolyte gel or paste (Driscoll et al., 2021). The EEG2/HdEEG arrays are chronically affixed to the skull surface with UV curable cement without the need for any skull burr holes. We validated this system by measuring normal and epileptiform EEG2 signals obtained with traditional metal epidural electrodes and MXene electrodes and comparing the spatiospectral distribution of preictal and epileptiform spectral density obtained with MXene HdEEG electrodes with those reported previously using epidural HdEEG arrays. Finally, we used MXene HdEEG arrays to determine the acute effects of mTBI on resting state spectral density and connectivity. Many mTBI patients have long-term neurological symptoms (Nelson et al., 2019) and elucidating the neurophysiological consequences of mTBI is a necessary step in developing effective mTBI therapies (Liang et al., 2022).

## 2. Materials and Methods

### 2.1 Animals

The protocols were approved by the [Author University] Animal Care and Use Committee. We used adult mice (20±3 weeks) in the DBA/2J background that heterozygously expressed a missense human epilepsy mutation in the GABAA receptor (Gabra1^+/A322D^) that that are an established model of epileptiform spike-wave discharges (Arain et al., 2012). Mice had unlimited access to food and water and were housed in a temperature- and humidity-controlled environment with a 12-hour lights-on/lights-off cycle with lights on at 0600 (Zeitgeber Time 0, ZT0).

### 2.2 Electrode array fabrication

We used free online software (EasyEDA) to design double sided PCBs for HdEEG arrays with 16 electrode pads (0.7 mm diameter rings with 0.2 mm holes) placed over the dorsal cortex and ground and reference electrodes placed over the midbrain (Fig 1A-B). EEG2 MXene arrays are identical to HdEEG arrays but only use the two recording electrodes over the left and right motor cortex (M3/M4, Fig 1A-B). Figure 1B-C shows the positioning hole used to align the arrays with bregma as well as the holes through which photocurable cement is applied to affix the arrays to the skull. The PCBs for the traditional epidural tungsten EEG2 arrays were similar to the MXene arrays except that the electrode pads were 1.05 mm diameter rings with 0.75 mm holes. For both arrays the wires connecting electrode pads were 150 μm wide. We designed the PCB to have pads for the attachment of either one four position (EEG2) or two ten position (HdEEG) rectangular connector sockets (Fig 1A-B) to connect the arrays to the EEG amplifier. The use of the four position EEG2 connector allows the array to interface with a commercial mouse EEG2 acquisition system (Pinnacle). Flexible PCBs using this design were manufactured on 0.2 mm polyimide by a commercial PCB prototyping company (PCBWay).

**Figure 1:**
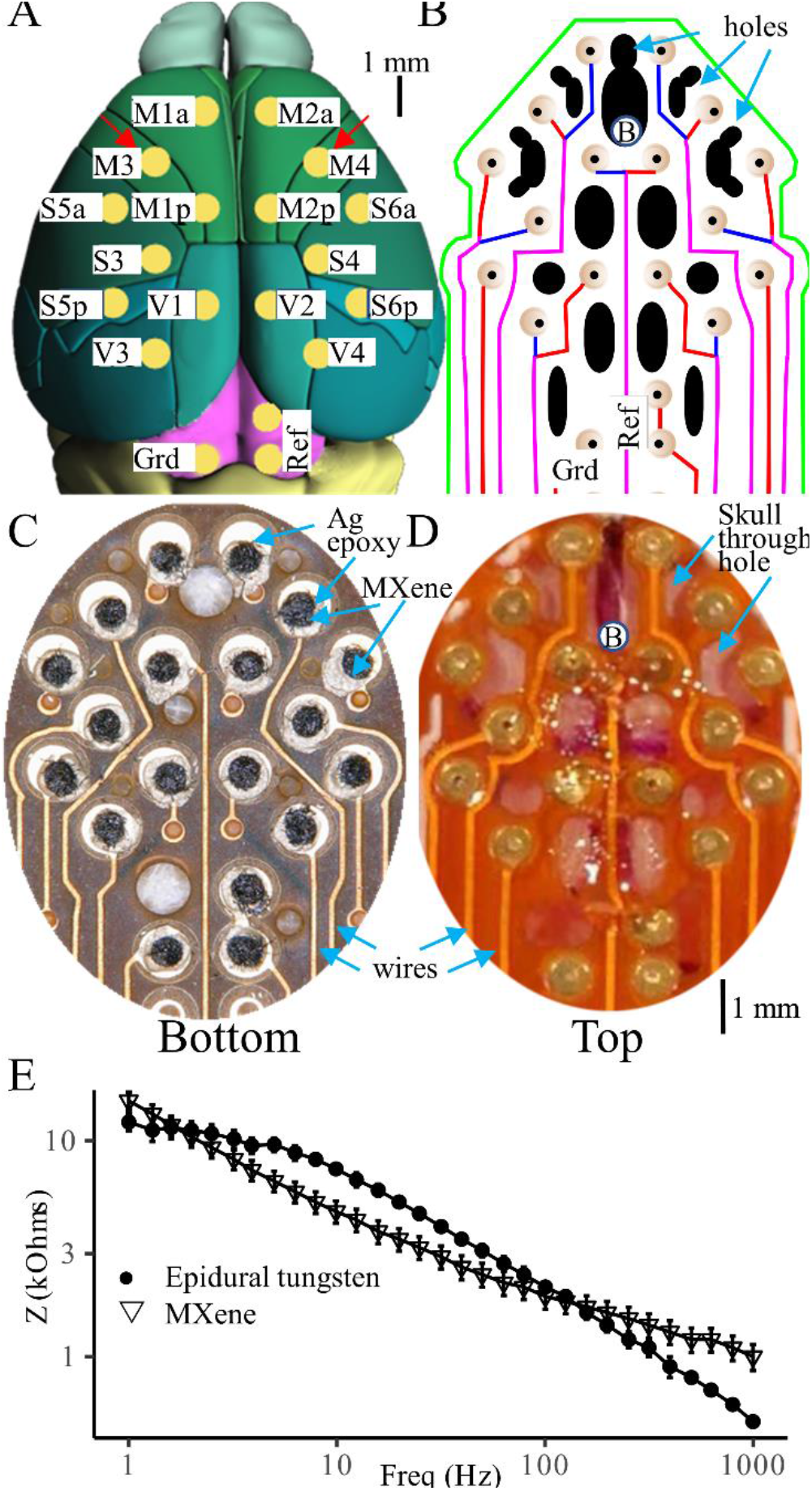
MXene HdEEG fabrication. A) Drawing of the mouse dorsal cortical surface showing the position of the 16 HdEEG channels (yellow) along with the ground (Grd) and reference (Ref) placed over the midbrain. The uppercase M, S, and V letters of the electrode names indicate placement over motor, somatosensory, and visual cortices. The lowercase a, and p letters in the M1/2 and S5/6 electrodes indicate anterior and posterior placement. Electrodes M3 and M4 indicated with red arrows are used for EEG2 recordings. Scale bar indicates 1 mm. B) Circuit diagram shows HdEEG PCB design. Electrode pads (beige) connected with wires (red = top PCB surface, blue = bottom PCB surface, magenta = separate wires on top and bottom surface) that ultimately terminate on connector pads. Black shapes indicate holes where UV-curable cement is placed. The ‘B’ placed at the bottom of the uppermost black hole indicates positioning over bregma that enables reproducible placement. C) Photograph of bottom of completed MXene HdEEG array. Silver epoxide is silver, MXene is dark gray, and wires are yellow. D) Intraoperative photograph of implanted MXene HdEEG array. Letter ‘B’ indicates bregma. E) Impedance spectra of epidural (●) and MXene (∇) electrodes measured in PBS solution (N = 5).

For the traditional epidural EEG2 arrays, five gold-plated electronic pin receptacle connectors (0.52 mm inner diameter, Mill-Max, 4428-0-43-15-04-14-10-0) were soldered to the electrode pads and then strengthened with nonconductive epoxy. The electrodes were five 3 mm tungsten rods (254 μm diameter, A-M Systems 717200) cut to 3 mm and ground to a 100 μm tip with a Dremel rotary tool.

For the MXene arrays, a polyimide template with holes overlying the electrode pads was made using a CO_2_ laser cutter. Ti_3_C_2_T_x_ MXene aqueous dispersions were provided by MuRata Manufacturing Co., Ltd. A nonwoven, hydroentangled cellulose polyester (60 to 40%) blend substrate was infused with the Ti_3_C_2_T_*x*_ MXene (20 mg/ml) and thoroughly dried. Circular electrode contacts were then cut out from the MXene-infused substrate using a biopsy punch to form circles (h 0.4 mm, d 500 μm). With the polyimide template placed over the PCB, the circular MXene contacts were applied to each electrode pad using conductive silver epoxy (CircuitWorks CW2400). Rectangular socket connectors were soldered to the PCB connector pads (four position connector for EEG2 cut from six position Samtec SFMC-103-T1-S-D; two ten position connectors for HdEEG cut from twenty position Samtec CLP-110-02-L-D-P-TR).

Samples of both the traditional tungsten electrodes and MXene electrodes were characterized by electrochemical impedance spectroscopy using a Gamry Reference 600 potentiostat. Measurements were conducted in room temperature 10 mM phosphate-buffered saline (PBS solution, pH 7.4) using a three-electrode configuration, with an Ag/AgCl reference electrode and a graphite rod counter electrode. Impedance spectra was collected at frequencies ranging from 1 Hz to 1 MHz with 10 mV rms AC voltage.

### 2.3 Surgery

Under isoflurane anesthesia, we made an incision in the dorsal scalp, cleaned, and dried the skull, and centered the positioning hole over bregma to ensure reproducible placement. For the epidural EEG2 arrays, the PCBs were attached to the skull with cyanoacrylate adhesive and dental cement. Then, a 29 Ga needle was inserted through the lumen of the electrode connectors to gently drill burr holes in underlying skull. The tungsten electrodes were then inserted in the connectors with their tips in the epidural space. The MXene HdEEG arrays were placed on the skull and a small amount of ultraviolet-curable adhesive (Visbella UF0008CR2P) was applied individually to each access hole while the nearby MXene electrode pads were held firmly against the skull to prevent the epoxy from seeping between the electrode and the skull. A UV light cured the epoxy.

### 2.4 EEG recording and analysis

The mice were acclimated to the EEG recording chamber for approximately two hours on postoperative days 0 and 1 after the implantation surgery. EEG2 and HdEEG recordings (approximately three hours) were performed using Nicolet V32 amplifier sampled at 250 or 500 Hz. For the experiments evaluating the durability of epidural and MXene EEG2 arrays, recordings were performed one, two and three weeks after surgery. Comparisons of awake background and preictal/ictal activity were made from recordings performed one week after surgery. Both types of recordings were performed between ZT3-ZT5. For the experiments that used MXene HdEEG arrays to determine the effects of mTBI on resting state spectral density and connectivity, the pre-mTBI/sham EEG was performed at ZT3 seven days after surgery. The mTBI/sham was done at ZT6 followed by a post-mTBI/sham recording that day (post TBI/sham 0) starting approximately ZT8. Additional post-mTBI/sham recordings were done at ZT3 on days 1 and 7 after the mTBI/sham. For all experiments, mice were killed by CO_2_ asphyxia on the last day of recording and their skulls and brains were examined to evaluate for the presence of visible injury.

A reviewer blinded to mouse identity and treatment identified EEG segments for analysis. Five segments (2s) of awake background EEG were chosen from each mouse containing predominantly low voltage and high frequency EEG activity (Louis et al., 2004) and without artifact or epileptiform discharges. SWDs were identified as high voltage rhythmic 6-8 Hz waveforms (Robinson and Gilmore, 1980; Sitnikova and Van Luijtelaar, 2007). Five SWD segments were chosen from each mouse from four seconds before- to two seconds after SWD onset (time 0, defined as first SWD spike).

EEG analysis was permed using the Fieldtrip MATLAB toolbox (Oostenveld et al., 2011). EEG was downsampled to 200 Hz, notch filtered at 60 Hz, and bandpass filtered between 0.5-55 Hz using a windowed-sinc finite impulse response filter (Widmann et al., 2015). Spectral power and cross spectral density were calculated using a two cycle Morlet transform from 2-55 Hz performed every 50 ms from 0.5 to 1.5 seconds of the awake background segments and from 0.5 to 1.5 seconds of the ictal segments. For the awake background, the spectral power and cross spectral density was averaged among the 50 ms segments. To compare results with a previous study, the pre-ictal analysis was done with Morlet transforms in the β spectral frequency (13-30 Hz) from −2.0 to −0.1 seconds. Awake background spectral density is given in relative spectral power, the spectral power at each integer frequency divided by the sum of the spectral power from 2-55 Hz. For preictal and ictal SWDs, normalized spectral power is reported in decibels (dB) and is calculated as:

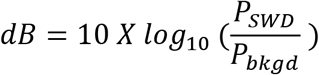

where P_SWD_ and P_bkgrd_ are the preictal/ictal and awake background spectral power, respectively. The weighted phase lag index (WPLI), a connectivity measure that is insensitive to volume conduction (Vinck et al., 2011), was calculated for each mouse from the cross spectral density. A bootstrap method was used to calculate the WPLI 95% confidence interval and the WPLI values with confidence intervals that did not overlap zero were considered suprathreshold (McGuinness et al., 2022). The network node degree, the number of suprathreshold connections, was determined at each electrode.

### 2.5 Overpressure blast mild traumatic brain injury

We used an established overpressure method for producing mTBI (Heldt et al., 2014; Guley et al., 2016, 2019). The mice were anesthetized with 2-4% isoflurane in 100% O_2_ throughout the procedure, first in an induction chamber and then via a nasal cannula while the mice were in the overpressure apparatus. After anesthesia induction, the mice were placed in a plastic holding tube which was then inserted in a second plastic tube that is situated on an x-y table that can precisely position the skull 4 cm from the barrel of a paint ball gun. The dorsum of the mouse skulls (at bregma) was positioned under 6 mm holes in the plastic tubes that align with the barrel and transmit the pressure blast to a precise location in the skull. The pressure of the paintball gun provided a single 40 psi pressure wave with a half-width duration of 7 ms; sham mice experienced the anesthesia and exposure to the apparatus without overpressure. After the sham/mTBI, we measured the righting time, a common immediate assessment of TBI severity (Kane et al., 2012; Gullotti et al., 2014). Immediately after the mTBI/sham, the mouse was placed on its side and the time until it stood on its paws was considered the righting time.

### 2.6 Statistical Analyses

Statistics were calculated using R for Windows version 4.1.3. Fabrication and surgery times and differences in mTBI/sham righting times were compared using two-tailed t-tests. Durability of epidural and MXene electrodes were compared using a log-rank test of the Kaplan-Meier estimates of survival. Perioperative deaths (within 7 days) were compared using the Fisher Exact test. Differences in spectral distribution of in the EEG2 awake and SWD data, and differences in spectral density and network node degree in the HdEEG experiments were calculated using cluster-based permutation (CBP) analyses. CBP analysis is a nonparametric statistical method (Maris and Oostenveld, 2007) that addresses the multiple comparison problem inherent in analyzing numerous electrodes and spectral frequencies by clustering nearby electrodes and frequencies based on either initial repeated measure (comparing pre-ictal and background spectral power) or independent (comparing pre-mTBI/sham, post sham, and post mTBI) T-tests. The summed T-score of the clusters were determined. The data sets were then randomly permuted 500 times and the percentile of the initial summed T-score versus the permuted T-scores is the probability that the spatiospectral distributions differ. Here, the CBP analyses were run with spatial clusters including electrodes within 2 mm and a critical cluster statistic of 0.025; P-values were appropriately adjusted to account for two tailed comparisons. In addition, a Bonferroni correction was made to the P-values when comparing the pre-mTBI/sham, post-sham, and post-mTBI mice.

## 3. Results

### 3.1 Array fabrication and implantation

Top and bottom views of completed MXene HdEEG arrays are shown in Figure 1C-D. Despite having 16, rather than two channels, the fabrication time of the MXene HdEEG-arrays (97±5 min) was significantly shorter than that of the epidural EEG2 arrays (179±57 min; p=0.004). Impedance spectroscopy demonstrated that the epidural and MXene electrodes had similar electrode/electrolyte impedances at frequencies between 1 and 1000 Hz (Fig 1E) with maximal difference of only 3.1 kΩ at 1 Hz.

Despite having a larger number of channels, the average duration of the implantation surgery for MXene HdEEG electrodes (55±4 min, N=7) was significantly shorter than that for epidural EEG2 arrays (73±4 min, N=12, P<0.001). The mice tolerated the implantation of both types of arrays. There were two peri-operative (within 7 days) deaths in the MXene HdEEG group and no deaths in the epidural EEG2 group (p=0.51).

### 3.2 EEG2 recordings with epidural and MXene electrodes

We directly compared epidural EEG2 recordings with the corresponding channels (M3/M4) of the HdEEG arrays in ambulating mice. Sample awake background EEG2 traces recorded by epidural and MXene electrodes are shown in figure 2A. Both electrode types provided excellent EEG quality. As expected, EEG2 from epidural electrodes was of higher voltage than MXene electrodes placed on the skull surface. However, MXene recordings depicted at higher gain (Fig 2A, bottom trace) showed similar waveforms without apparent noise interference. Spectrograms (Fig 2B) show that awake background recordings by epidural and MXene EEG2 systems exhibit similar distributions of relative spectral density although the MXene electrodes exhibit slightly increased 2-3 Hz and 20-30 Hz density while the epidural electrodes exhibit slightly increased 7-14 Hz density (P=0.02).

**Figure 2:**
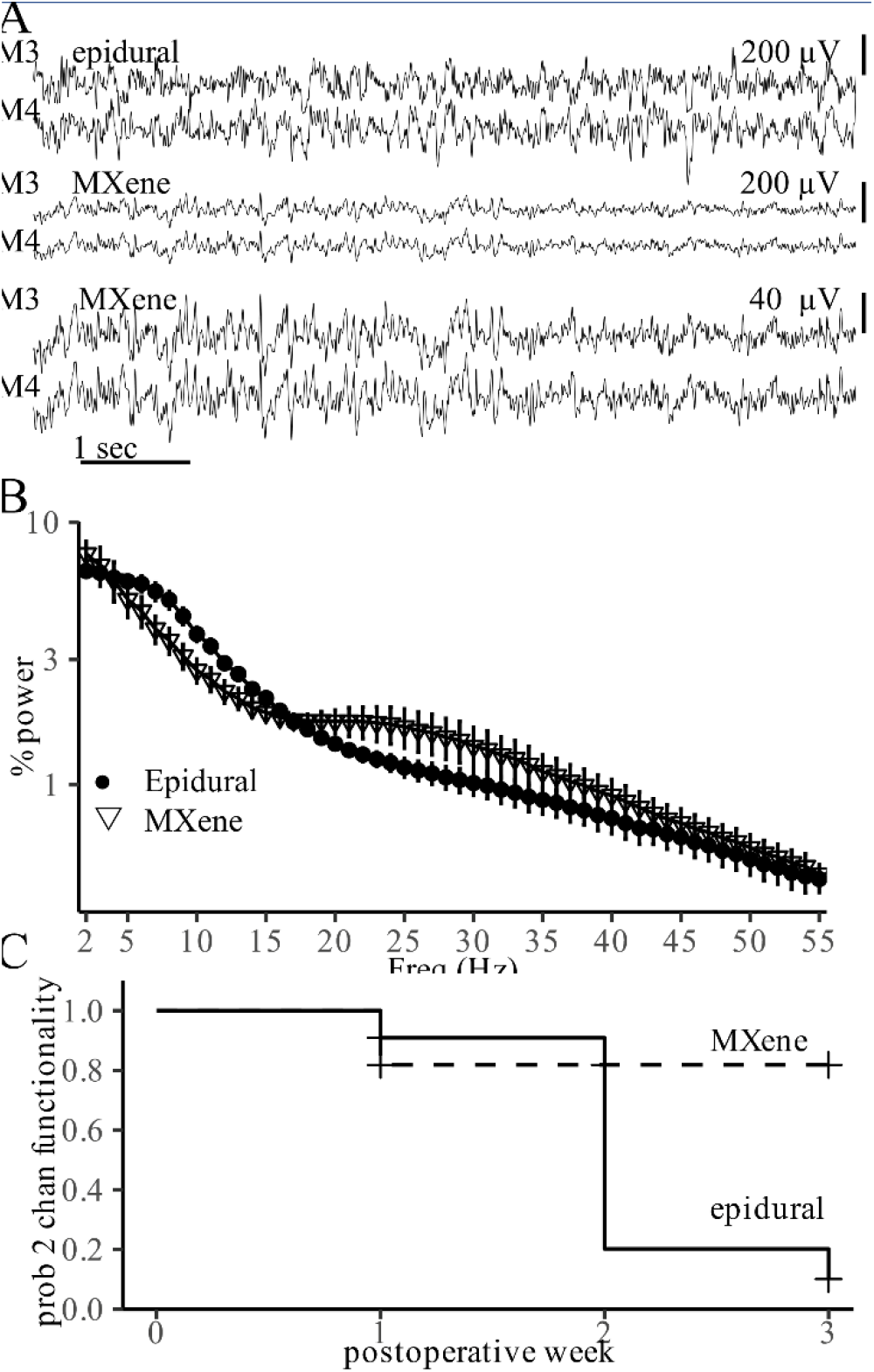
Epidural and MXene EEG2 recordings of awake background. A) Sample EEG2 recordings (channels M3/M4) of awake background obtained by epidural electrodes (top) and MXene electrodes (middle and bottom). Scale bars indicate sensitivity of each trace. B) Mean wake background relative spectral power ± SEM obtained with epidural (●, N = 10) and MXene (∇, N = 6) electrodes (P=0.02). C) Probability of both EEG2 channels functioning after implantation with epidural (solid line) and MXene (dashed line) electrodes (N=11, P=0.02, cluster 8-11 Hz)

We determined the durability of the epidural and MXene electrodes in providing quality EEG2 in both channels (Fig 2C). While the 100 μm diameter epidural electrodes allow smaller skull burr holes than 1 mm epidural screw electrodes, they are less durable than large screws. Consequently, 82% of the epidural preparations lost one channel within three weeks of implantation. In contrast, MXene electrodes provided significantly better reliability than the 100 μm epidural electrodes (P = 0.02) with only 18% of the mice losing one of the EEG2 channels at three weeks.

Next, we compared the EEG2 recordings of epileptiform SWDs obtained with epidural and MXene electrodes. SWDs obtained with both types of electrodes showed the typical spike, positive transient, and wave components reported previously in rodent absence seizures (Robinson and Gilmore, 1980; Sitnikova and Van Luijtelaar, 2007). Spectrograms of the SWD background-normalized spectral density are shown in Figure 3B. Both epidural and MXene SWD recordings exhibited a broad peak of normalized spectral density from 6-20 Hz reflecting the 6-8 Hz primary SWD frequency along with the first and second harmonics. Because skull surface MXene electrodes recorded awake background EEG at lower voltage than epidural electrodes, MXene electrodes exhibited a higher SWD background-normalized spectral density (P = 0.02)

**Figure 3:**
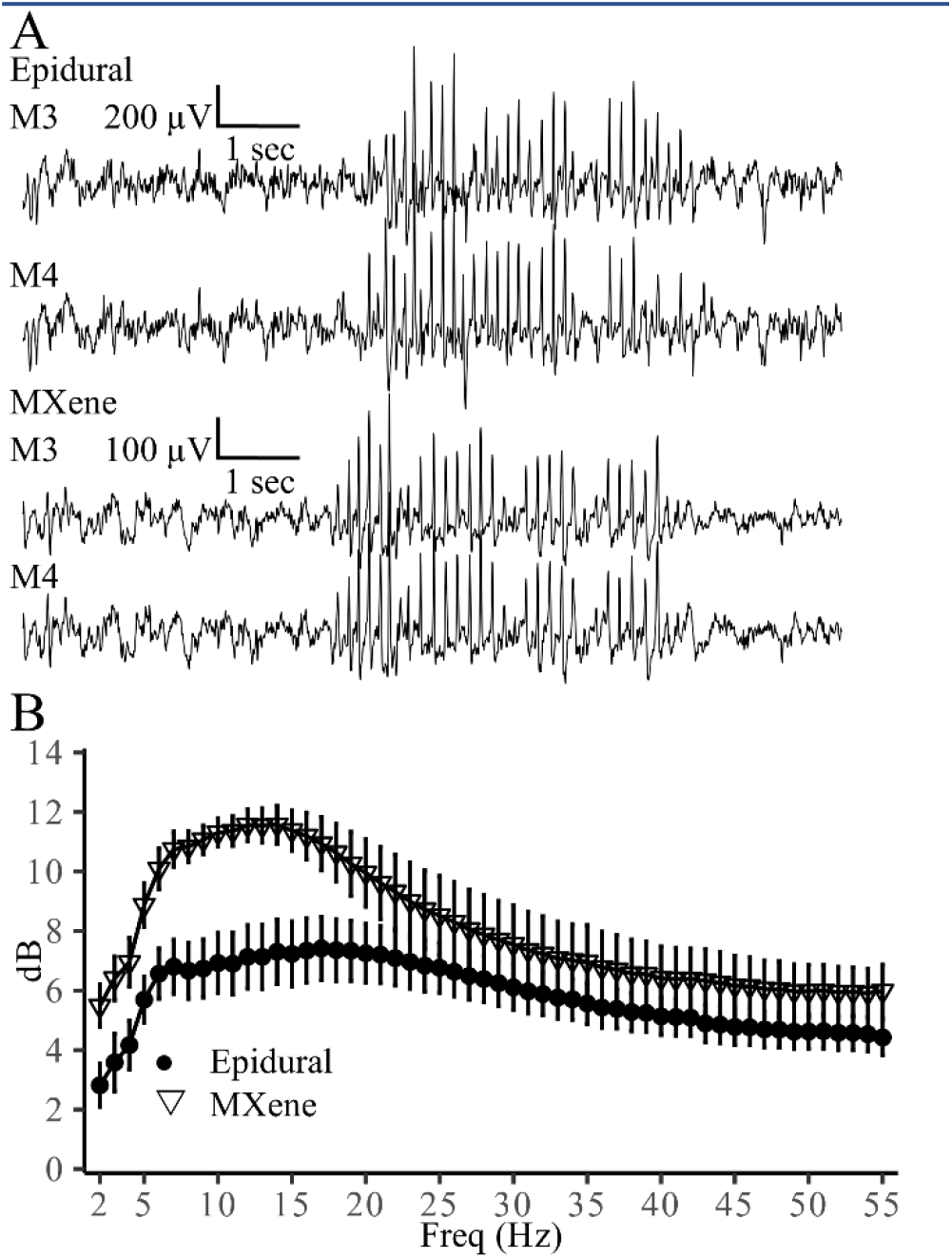
Epidural and MXene EEG2 recordings of SWDs. A) Sample EEG2 recordings (channels M3/M4) of spike-wave discharge obtained with epidural electrodes (top), or MXene skull surface electrodes (bottom). Scale bars indicate sensitivity. B) Spike wave discharge mean background-normalized spectral power ± SEM obtained with epidural (●, N=9) and MXene (∇, N=6) electrodes (P=0.02, cluster 5-16 Hz).

### 3.3 MXene HdEEG SWD recording

We next recorded HdEEG using the MXene electrodes. Like the M3/M4 electrodes (Fig 2C), most of the additional MXene HdEEG electrodes, electrodes - with the exception of the S5p and S6p electrodes - provided excellent EEG signal for at least three weeks (not shown). However, the S5p and S6p electrodes in the lateral posterior region were more likely than the other HdEEG to become dislodged early after implantation (week 1, with probabilities of 0.38 and 0.31, respectively).

Previous studies used epidural HdEEG electrodes to map the spatial distribution of SWDs (Ding et al., 2019) and epidural HdEEG or single electrodes to identify preictal increases in β frequency spectral density occurring prior to SWD onset (Sorokin et al., 2016; Ding et al., 2019). Therefore, we recorded SWDs with MXene HdEEG electrodes and compared their spatial distribution and pre-ictal β frequency spectral density with those obtained in the prior studies using epidural electrodes. An HdEEG SWD tracing is depicted in Figure 4A and a topology plot of the mean 6-8 Hz spectral power of the SWD onset (0.5-1.5 s after first spike) is shown in Figure 4B. As seen previously with epidural HdEEG electrodes (Ding et al., 2019), the characteristic 6-8 Hz spikes and waves are recorded in all electrodes over the dorsal cortex, but with a prominence in the frontal electrodes over the motor cortex.

**Figure 4:**
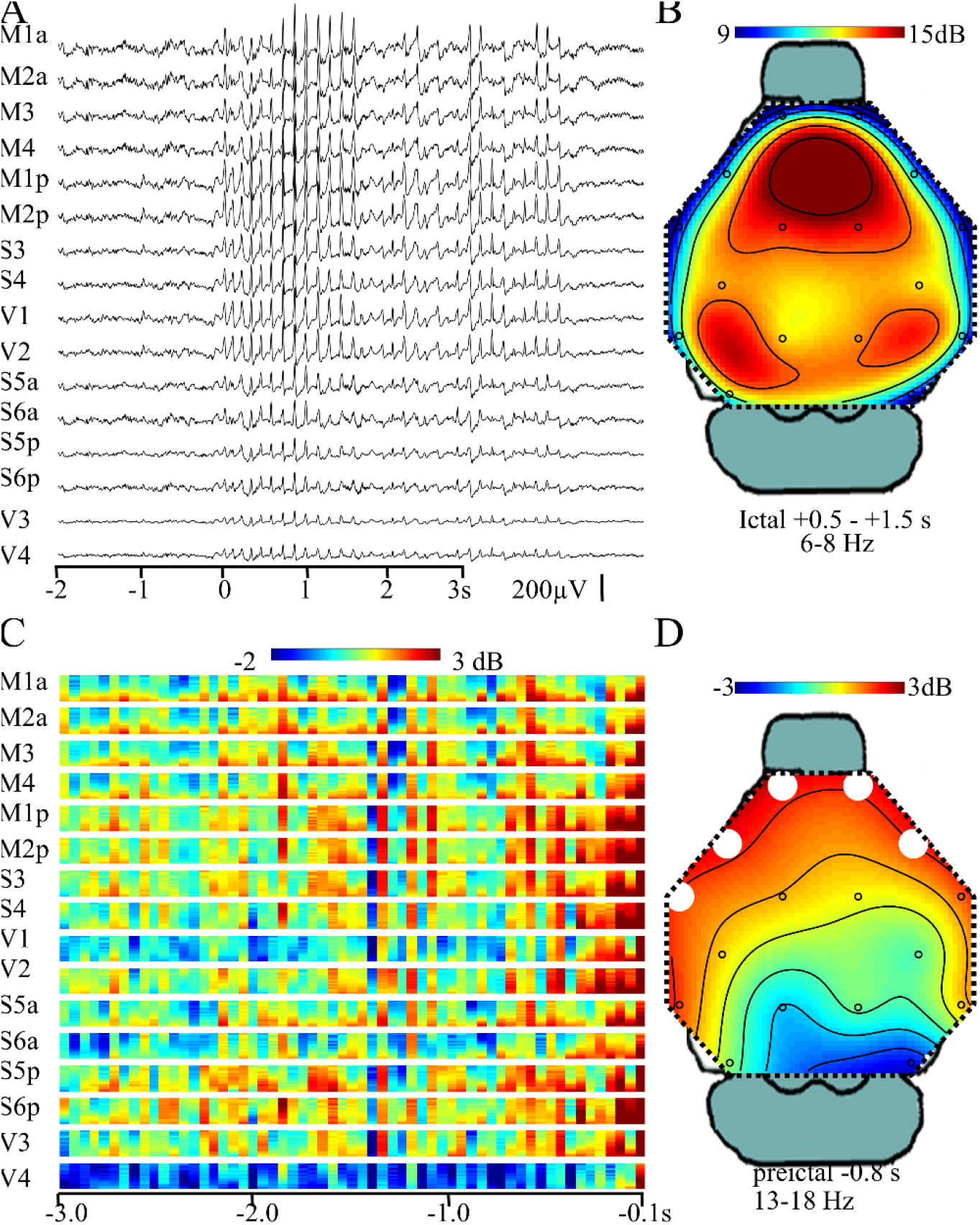
MXene HdEEG SWD. A) Sample HdEEG recording of a SWD obtained with MXene skull surface HdEEG. SWD onset (t_0_) is the time of the first spike. B) SWD mean background-normalized spectral density (6-8 Hz, 0.5-1.5s) topology plot (N = 12). C) Mean pre-ictal beta frequency (13-30 Hz, each y axis) background-normalized spectral power at each HdEEG channel plotted from 3.0 seconds to 0.1 second prior to SWD onset. Times −1.0 to −0.1 seconds statistically compared with awake background 13-30 Hz power (N=12, P=0.002). D) Topology plot of mean 13-18 Hz spectral power at 0.8 seconds prior to SWD onset. White circles indicate electrode cluster found in the CBP analysis.

Figure 4C depicts the mean preictal (3.0 to 0.1 s before first spike) β frequency (13-30 Hz) background-normalized spectral density and demonstrates an increase in spectral power first over the motor and somatosensory cortices shortly after one second prior to seizure onset followed by increased beta power over the visual cortex. CBP testing (13-30 Hz) revealed a significant difference between the pre-ictal period (between −1.0 and −0.1s) and awake background with clusters between −0.8 and −0.1 seconds and 13-18 Hz (Fig 4D, P = 0.002).

### 3.4 Effect of mTBI on awake HdEEG spectral power

One week after MXene HdEEG implantation, mice underwent a two-hour baseline recording followed by either mTBI or sham. The mTBI produced no difference in visible behavior or righting times (mTBI 107±19s, sham 113±17s). Subsequent HdEEG recordings at two hours, one day and seven days after mTBI/sham demonstrated that mTBI did not disrupt MXene electrode recording quality. Postmortem examination of the brains found no visible evidence of injury in either group (not shown).

Figure 5A depicts anterior (M2a) and posterior (V2) HdEEG time-series EEG obtained prior to mTBI/sham (pre) as well as two hours after sham or mTBI and Figure 5B demonstrates the mean pre/sham/mTBI relative spectral density and their confidence intervals. Grids depicting the mean differences in the relative spectral density between pre-mTBI/sham and two hours post sham (left) or post-mTBI (right) for all HdEEG channels excluding S5p and S6p (due to the diminished durability of these electrodes) are shown in figure 5C. Even though mTBI did not alter visible behavior or righting times, it did significantly alter the distribution of spectral density (P = 0.001). Examination of the grids (Fig 5C) and topographic plots (Fig 5D) demonstrates that mTBI redistributes relative spectral density from low frequencies to β-frequencies (12-30 Hz) predominantly in the central and posterior regions.

**Figure 5:**
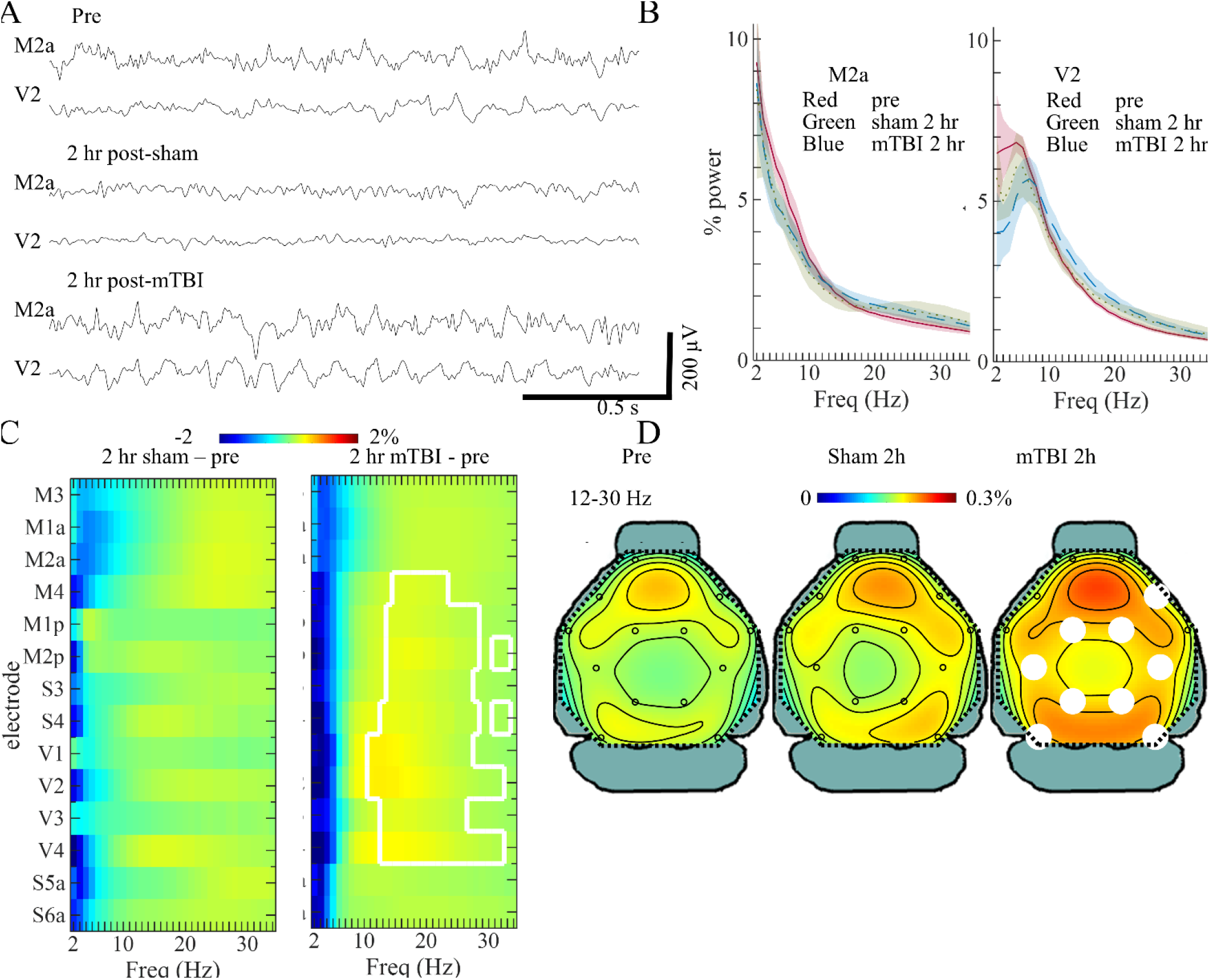
Acute mTBI effects on awake spectral density. A) Sample anterior channel (M2a) and posterior channel (V2) MXene HdEEG recordings of awake background prior to mTBI (top) and two hours after sham (middle) or mTBI (bottom). B) Mean relative spectral density from anterior (left) and posterior (right) HdEEG channels prior to mTBI/sham (red, N=13), or two hours after sham (green, N=6) or mTBI (blue, N=8). Shaded regions represent 95% confidence intervals. C) Grids depict mean differences in relative spectral power between pre mTBI/sham and two hours post sham (left) or mTBI. CBP testing found a significant difference in the spatiospectral distribution between pre-mTBI/sham and mTBI (P=0.001, Bonferroni corrected). Electrodes/frequencies enclosed by white lines indicate CBP clusters. D) Topology plots depict relative spectral density of pre-mTBI/sham (left), sham (middle), and mTBI (right) 12-30 Hz (bottom row). White circles indicate channel clusters from CBP testing.

The effects of mTBI on relative spectral density were not present one or seven days after the mTBI and there were no significant changes in network node degree at any of the time points (not shown).

## 4. Discussion

This study demonstrated high quality skull surface EEG2 and HdEEG recordings using open-source MXene-based EEG arrays. The signals recorded with MXene electrodes were comparable to those obtained with epidural electrodes. MXene HdEEG recordings revealed that mTBI produced acute changes in spatiospectral distribution of spectral power, despite the absence of visible behavioral differences. These results demonstrate that that the MXene skull surface electrode array is a novel, inexpensive tool that allows a minimally invasive mechanism to study mouse neurophysiology.

### 4.1 Comparison of MXene electrodes with current techniques

Compared with current epidural EEG2 and HdEEG methods (Wasilczuk et al., 2016; Ding et al., 2019), this open source method of MXene electrode skull surface EEG eliminates the need for drilling skull burr holes. This advance represents a major refinement in animal care. Eliminating skull burr holes also mitigates potential confounding factors in rodent physiology experiments, especially those designed to study the effects of TBI or those investigating physiology of young mice. MXene EEG array fabrication is more efficient than the epidural open-source techniques given that its layout and wiring are achieved using PCBs and its electrode pads are easily coated with MXene. Importantly, surgical implantation of MXene arrays is easier and more efficient than epidural electrodes since burr holes are not required and may thus be performed by laboratory personnel with less experience than those who typically perform intracranial surgeries.

Nanofabricated 16 and 32 channel skull surface HdEEG arrays are commercially available (Neuronexus) and are substantially thinner (15 μm) than the ones produced by our open-source method. However, although the electrodes of the commercial arrays lie on the skull surface, chronic placement of this array still requires three skull penetrating anchoring screws (Jonak et al., 2018) and thus is problematic for mTBI studies and experiments using young mice with thin skulls. Another substantial advantage our open-source MXene HdEEG system compared with commercially available arrays is reduced cost (US$6.64 vs US$778). Although a technique has been reported allowing retrieval and reuse of the commercial HdEEG up to six times (Jonak et al., 2018), the cost per use of the commercial array is still substantially greater than that of the MXene HdEEG system. Therefore, our open-source method will be especially beneficial for investigators needing to screen many subjects with chronic EEG.

Although most MXene electrodes provided quality EEG signal for at least three weeks after implantation, the S5p/S6p lateral-posterior region were more susceptible to displacement. In the current design (Fig 1), these two electrodes lack an adjacent hole through which UV-curable cement is applied. Future studies using a modified MXene HdEEG design with a cement attachment site near the lateral-posterior electrodes will determine the long-term usability of these electrodes. Because the MXene HdEEG electrode arrays can be removed by gently scraping the cement from the skull surface, these experiments will also determine the feasibility of replacing the arrays if electrodes become dislodged.

### 4.2 mTBI acutely alters the distribution of resting state relative spectral power

Although previous studies have used epidural EEG2/limited-channel EEG to measure the effects of mTBI in mice (Lim et al., 2013; Modarres et al., 2017; Ichkova et al., 2020; MacMullin et al., 2020), this is the first report, to our knowledge, that measured the effects of mTBI on resting state EEG using non-penetrating electrodes or with HdEEG. We found that although mTBI did not change righting time compared with sham exposure and there were no visible behavior differences between mTBI and sham mice, mTBI did acutely change relative spectral power with the most notable increases in β spectral frequency in the central and posterior electrodes. Thus, the significant change in the distribution of relative spectral power could serve as a biomarker of injury. Future experiments that perform MXene HdEEG in conjunction with formal behavioral testing and postmortem immunohistological analysis with markers of microscopic injury will determine if the electrophysiological changes predict behavioral dysfunction or tissue injury.

Mild TBI increases the risk for developing epilepsy (Annegers et al., 1998; Christensen et al., 2009; Karlander et al., 2021) and early seizures, those that occur within seven days of the injury, may increase that risk (Lowenstein, 2009). Because epileptiform SWDs are preceded by increased β-spectral power (Ding et al., 2019; Sorokin et al., 2016, and Figure 4) it is possible that mTBI’s acute effect on increasing resting-state β-spectral power (Figure 5) may raise the risk of early seizures. Future studies will determine the association of mTBI, β-spectral power, and early seizures.

### 4.3 Conclusions

In conclusion this study determined that open source MXene EEG2 and HdEEG electrodes represent an inexpensive, convenient method for recording mouse EEG without skull disruption and represent a refinement in animal care. Using MXene HdEEG recordings, we determined that mTBI acutely altered the spatiospectral distribution of relative spectral power with increased β spectral power in the central and posterior electrodes, a possible biomarker of mTBI-associated cerebral dysfunction.

## Notes

### Competing Interest Statement

The authors have declared no competing interest.

